# TRIB1 regulates tumour growth via controlling tumour-associated macrophage phenotypes and is associated with breast cancer survival and treatment response

**DOI:** 10.1101/2021.06.07.446596

**Authors:** Taewoo Kim, Jessica Johnston, Francisco J. C. Felipe, Stephen Hamby, Sonia Castillo-Lluva, The Cardiogenics Consortium, Alison H Goodall, Guillermo Velasco, Alberto Ocana, Munitta Muthana, Endre Kiss-Toth

## Abstract

Molecular mechanisms that regulate tumour-associated macrophage (TAM) phenotype and function are incompletely understood. Here, we show that the pseudokinase TRIB1 is highly expressed by TAMs in breast cancer and that its expression correlates with response to chemotherapy and patient survival. We used immune-competent murine models of breast cancer to characterise the consequences of altered (reduced or elevated) myeloid *Trib1* expression on tumour growth and composition of stromal immune cells. We found that both overexpression and knockout of myeloid *Trib1* promote tumour growth, albeit through distinct molecular mechanisms. Myeloid *Trib1* deficiency resulted in an early accelearation of tumour growth, paired with a selective reduction in perivascular macrophage numbers *in vivo* and enhanced oncogenic cytokine expression *in vitro*. In contrast, elevated levels of *Trib1* in myeloid cells led to an increase in mammary tumour volume at late stages, together with a reduction of NOS2 expressing macrophages and an overall reduction of these cells in hypoxic tumour regions. In addition, we show that myeloid *Trib1* is a previously unknown, negative regulator of the anti-tumour cytokine IL-15 and that increased expression of myeloid *Trib1* leads to reduced IL-15 levels in mammary tumours, with a consequent reduction in the number of T-cells, that are key to anti-tumour immune responses.

Together, these results define the different roles of TRIB1 in human breast cancer and provide a mechanistic understanding for the importance of myeloid TRIB1 expression levels in the development of this disease.

**Significance:** *TRIB1* expression is strongly associated with response to chemotherapy in breast cancer patients with aggressive tumours. This protein is also highly expressed by tumour-associated macrophages. Thus, we used myeloid-specific alterations of *Trib1* expression in mice (*Trib1^mKO^* and *Trib1^mTg^*), and characterised consequent changes in the growth rate and tumour microenvironment of mammary tumours. Both *Trib1^mKO^* and *Trib1^mTg^* enhanced tumour growth, but at different stages of tumour growth and via distinct mechanisms. *Trib1^mKO^* significantly increased the expression of oncogenic cytokines, such as IL6, IL10, CCL20, PD-L1, and VEGF. In contrast, *Trib1^mTg^* accelerated at the later stage of tumour growth via inhibition of hypoxic TAMs in the TME, as well as by reduced IL-15 expression thus leading to impaired *naïve* and cytotoxic T cell infiltration. These data define TRIB1 as a potential novel marker of therapeutic responses in breast cancer, as well as a key mechanistic regulator of the anti-tumour cytokine, IL-15 in myeloid cells.

## Introduction

Breast cancer (BC), the leading cause of cancer death in females (Torre et al., 2015), is initiated by the formation of a tumour niche where cancer-initiating cells or breast cancer stem cells recruit healthy, non-transformed cells (Feng et al., 2018, Sin and Lim, 2017). These cells are re-educated by stimuli from cancer cells to promote the expression of oncogenic cytokines and growth factors (Chen et al., 2015). Tumour-associated macrophages (TAMs) are one of the most abundant cell types that can comprise up to 50% of the tumour microenvironment (TME) and facilitate tumour initiation and development (Guo et al., 2016). A number of published studies reported high infiltration of TAMs in breast cancer, correlating with poor prognosis and clinical outcomes (Zhang et al., 2013, Yuan et al., 2014, Zhao et al., 2017, Qiu et al., 2018). Triple-negative breast cancer (TNBC) tumours were shown to have a higher number of CD68^+^ macrophages compared to other subgroups (Volodko et al., 2019). The plasticity of infiltrated TAMs is influenced by environmental signals and these cells can be functionally classified into M1 (pro-inflammatory) and M2 (anti-inflammatory) cells, as two extremes (Qiu et al., 2018). Though TAMs are able to express markers of either polarisation phenotype, pro-inflammatory macrophages are generally observed upon entering to the tumour site (Yuan et al., 2015), but M1-like macrophages stimulated by the type 1 T helper cell (Th1) cytokines are known to exhibit anti-tumour capacity by generating anti-tumour cytokines (such as TNF, IL-2, and IL-12) and reactive nitrogen and oxygen intermediates (Choudhari et al., 2013, Quatromoni and Eruslanov, 2012, Qiu et al., 2018). Most of TAMs are polarised to have a M2-like phenotype after infiltration which disrupts the clinical outcomes and patient survival (Macciò et al., 2020) and produce anti-inflammatory cytokines (such as IL-4) and growth factors to inhibit immune response and promote proliferation (Lin et al., 2019).

Hypoxia promotes the dephosphorylation of chemoattractant receptors and inhibits migrating stimulating factors that trap macrophages and TAMs in the hypoxic area and is associated with aggressive breast tumour phenotypes (Lin et al., 2019, Obeid et al., 2013). The entrapped hypoxic TAMs facilitate tumour vascularisation and immune suppression by expressing angiogenic molecules and immunosuppressive factors (Obeid et al., 2013, Henze and Mazzone, 2016).

The pseudokinase Tribbles-1 (TRIB1) is highly expressed in the macrophage lineage and has been shown to regulate macrophage polarisation (Richmond and Keeshan, 2019, Johnston et al., 2019, Niespolo et al., 2020) and *Trib1*-deficient mice were shown to lack anti-inflammatory macrophages (Satoh et al., 2013, Ye et al., 2012, Lee et al., 2014).

Using Bayesian network inference modelling, TRIB1 expression was shown to be correlated with the levels of NF-κB and IL-8 in breast cancer, and was also considered as a potential biomarker for clinical outcomes (Gendelman et al., 2017). However, unlike recent papers reporting an oncogenic role of TRIB1 in prostate cancer via regulating macrophage infiltration and inducing M2-like polarisation (Liu et al., 2019), to date no study has examined the TAM-specific tumoural capacity depending on *Trib1* and how these influence tumour development. Therefore, we examined mammary tumour development in mice where levels of *myeloid-Trib1* (*mTrib1*) have been regulated genetically. Based on our analyses, we demonstrate that both overexpression and knockout of *mTrib1* promote tumour growth, albeit through distinct molecular mechanisms and at different stages of tumour growth, providing a novel mechanistic insight into the functional consequences of TAM phenotypes.

## Results

### *TRIB1* is highly expressed in tumour-associated macrophages and its expression correlates with response to chemotherapy and patient survival in breast cancer

Whilst *TRIB1* is oncogenic in several cancer settings, its potential importance in BC pathogenesis and response to therapy are largely unknown. To explore the potential role TRIB1 may play in BC, we have analysed the correlation between TRIB1 mutations and BC survival in a dataset of 6697 patients (Pongor et al., 2015), using the G-2-O algorithm, as described before (Cimas et al., 2020). This method allows to establish the association between prognosis of a specific transcriptomic signature linked with a mutation and patient survival, by establishing two cohorts, the wild type and the mutant, both compared using a Cox survival analysis. Our analysis showed a highly significant reduction in patient survival in tumours with TRIB1 mutations (Figure 1A, HR 0.56 CI 0.50-0.62; lograk p=4.1×10^-26^).

**Figure 1.**
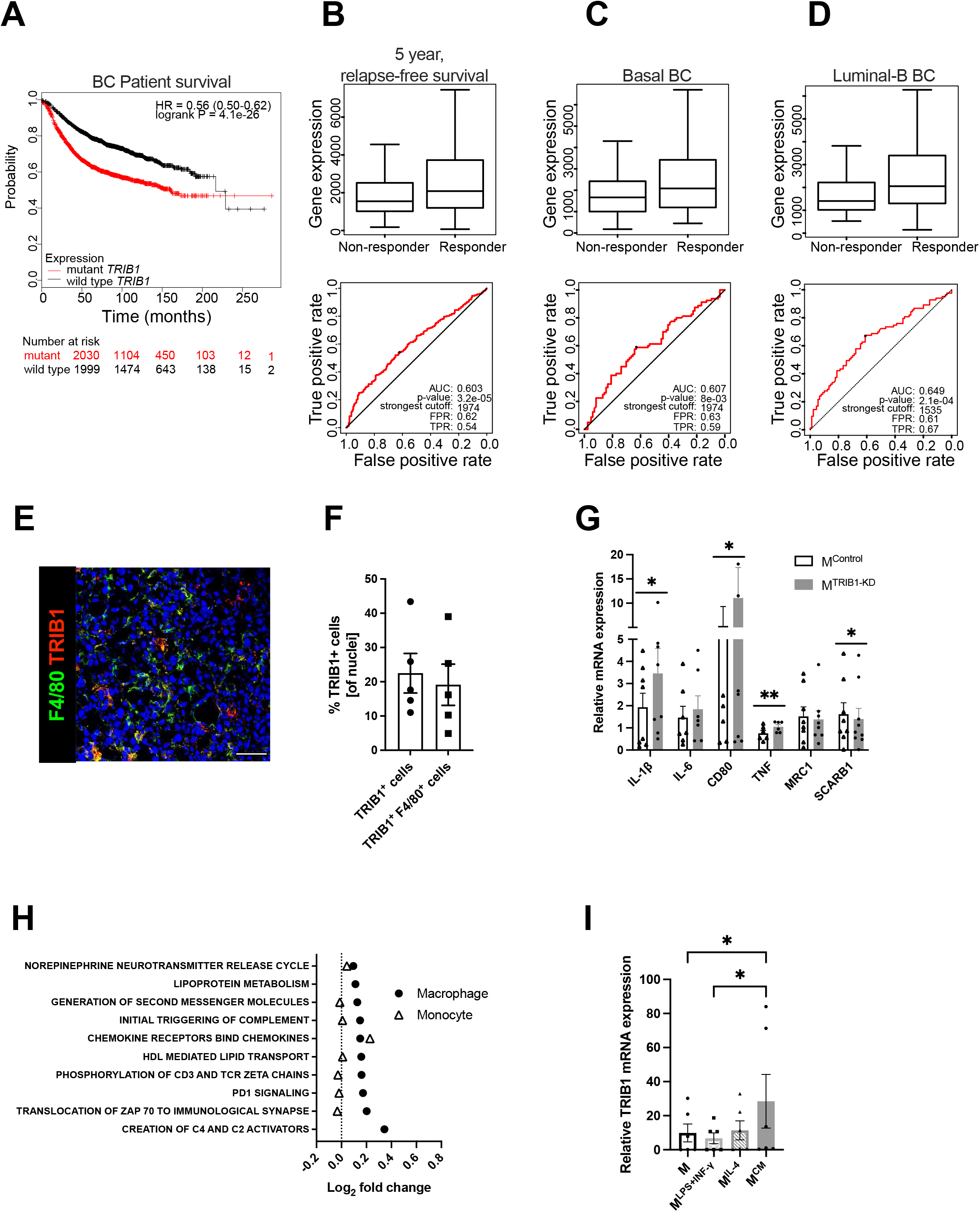
TRIB1 is associated with overall BC survival and 5-years relapse-free cancer survival after chemotherapy and is a highly expressed protein in Tumour-Associated Macrophages. (**A**) TRIB1 mutation-harbouring tumours are associated with a poor long-term BC survival. Kapplan-Maier survival plot, Hazard Ratio (HR) and log rank p value are displayed. False discovery rate (FDR). Red line represent patients harbouring the transcriptomic fingerprint of the mutation compared with those without in the dark line (**B)** *TRIB1* expression in 5 year relapse-free BC survival after anthracycline-based chemotherapy. Note that enhanced TRIB1 expression correlates with a higher 5 year relapse-free BC survival after anthracycline-based chemotherapy. Receiver Operating characteristic (ROC) curves are shown. Area under the curve (AUC), p value, false positive rate (FPR) and true positive rate (TPR). All breast cancer patients, basal-like and lumina B subtypes (**B, C, D, respectively)** Representative image of TRIB1 (red) and F4/80 (green) fluorescence staining of primary murine breast tumours from wild-type (C57/BL6) animals (Scale: 50μm). **(F)** Quantification of TRIB1 expressing cells and F4/80+ cells in the TME from “E” relative to total cell counts (n=5 mice/group). Data presented as mean±SEM. Cells in 5 random fields of views for each animal were manually quantified using ImageJ. **(G)** Human MDMs were transfected with either control or *TRIB1* siRNA for 48 hours. RNA levels of *IL-1β, IL-6, CD80, TNF, MRC1, and SCARB1* were quantified 48 hours after *TRIB1* siRNA transfection (M^TRIB1-KD^). Results of paired t-tests are presented; mean±SEM is plotted; *p<0.05 **p<0.01 (n=6-9 donor/group). **(H)** Pathway analysis of TRIB1 associated genes in monocytes and MDMs in participants of the Cardiogenics Transcriptomic Study. MDM (n=596) and monocytes (n = 758) were ranked according to *TRIB1* RNA levels and genes that were differentially expressed between the top vs. bottom 25% of the rankings were analysed with QuSAGE. The 10 most significantly enriched pathways in MDMs are presented. **(I)** *TRIB1* RNA levels of human MDMs unstimulated (M) and stimulated with LPS and INF-γ (M^LPS+INF-γ^), IL-4 (M^IL-4^), and CM (M^CM^) for 24 hours. Results of Friedman test and Dunn’s multiple comparison tests are presented; mean±SEM is plotted; *p<0.05 (n=6 donors/group).

Next, we evaluated whether the expression of TRIB1 correlated with relapse-free survival in BC patients by interrogating a dataset including 1329 patients with RFS information at 5 years (Fekete and Gyorffy, 2019). We analyzed different types of therapy, from endocrine treatment (tamoxifen and aromatase inhibitor) to specific anti-HER2 inhibitors (trastuzumab or lapatinib), and several chemotherapy treatments, including taxanes, anthracyclines, Ixabipelone, CMF (cyclophosphamide, methotrexate, fluorouracil), FAC (fluorouracil, adriamycin, citroxan) and FEC (fluorouracil, epirubicin, cyclophosphamide); and found that TRIB1 expression specifically correlates with 5 year relapse-free survival in anthracycline-based chemotherapy in overall groups of BC patients (Figure 1B) and specifically in Basal (Figure 1C) and Luminal-B (Figure 1D) BC subtypes. Once identified the potential role of TRIB1 in breast cancer outcome and response to therapy, we aimed to characterise TRIB1 protein expression within mammary tumours, using a murine model where BC growth was induced in immune-competent, C57BL/6 mice with an orthotropic injection of a murine, Basal-B BC cell line, E0771 (Le Naour et al., 2020) (Figure 1E). This analysis revealed that up-to 40% of cells expressed high levels of TRIB1 protein and that about 85% of these cells were also positive for the macrophage marker F4/80 (Figure 1F), suggesting a potential tumourigenesis regulating role for *mTrib1* in BC.

In order to gain an initial mechanistic insight into TRIB1-dependent alterations, relevant to tumour-biology, we used human monocyte-derived macrophages (MDMs) isolated from healthy human blood and transfected them with *TRIB1* siRNA to reduce the expression (M^TRIB1-KD^) to assess the expression of a range of genes, known to play an important role in TAM function. This analysis revealed that the knockdown of *TRIB1* in MDMs (for knockdown efficiency, see Figure 3M) enhances their pro-inflammatory phenotype with a significant increase of *IL-1β* (p<0.05), *CD80* (p<0.05), and *TNF* (p<0.01) levels and reduced *SCARB1* expression (p<0.05) (Figure 1G), in line with changes observed in MDMs stimulated towards an inflammatory phenotype (M^LPS+INF-γ^) as well as reports of transcriptomic changes in TAMs (Orecchioni et al., 2019) (Supplementary figure 1) (for knockdown efficiency, see Figure 3M). Next, we carried out a gene enrichment analysis with QuSAGE (Yaari et al., 2013, Turner et al., 2015, Meng et al., 2019) to identify biological pathways associated with altered *TRIB1* expression in human monocytes (n=758) and MDMs (n=596), using data from the Cardiogenics Transcriptomic Study (Heinig et al., 2010, Schunkert et al., 2011, Rotival et al., 2011). Comparison of the 10 most significantly enriched pathways in MDMs (Macrophage) vs. monocytes revealed that most of these pathways were only enriched in macrophages, confirming the distinct regulatory impact of *TRIB1* between these cell types (Figure 1H, Supplementary table 2). From those pathways significantly associated with *TRIB1* levels in macrophages, such as creation of C4 activators, translocation of ZAP 70 to immunological synapse, PD1 signalling, and phosphorylation of CD3 and zeta chains have previously been shown to be involved in the promotion of tumour growth and regulate T-cell activation and polarisation (Blanchard et al., 2002, Roumenina et al., 2019). In addition, increased PD1 signalling has been reported to increase macrophage proliferation and activation, and inhibit phagocytosis and tumour immunity in TAMs (Gordon et al., 2017, Hartley et al., 2018). Finally, treatment of MDMs with cancer cell-conditioned medium (CM) also showed a significant overexpression of *TRIB1* in these cells (M^CM^), compared to control (M^UN^) and M^LPS+INF-γ^ cells (p<0.05) (Figure 1I), suggesting a 2-way regulation of TRIB1 expression between BC cells and tumour macrophages.

### Mammary tumour growth is accelerated by *Trib1* alteration

Based on the above evidence of potential association between TRIB1 and BC, we hypothesised that myeloid TRIB1 expression may influence the aggressiveness of BC and thus modulation of *Trib1* expression in these cells would alter tumour growth. To test this, we used myeloid-specific *Trib1* overexpressing (*Trib1^mTg^*) and knockout (*Trib1^mKO^*) animals *vs*. their littermate controls, on an immune-competent, C57BL/6 background. We have recently reported details of the development and initial characterisation of these mouse lines (Johnston et al., 2019). We modelled BC growth, using these myeloid-specific mouse lines and a murine breast cancer cell line, E0771, that has recently been reported to be a luminal-B subtype (Le Naour et al., 2020), thus closely resembling the human BC subtype, where TRIB1 expression levels closely correlated with response to chemotherapy (Figure 1D). Mammary fat pads of 8 week-old mice were injected with E0771 cells, and the rate of tumour growth measured (Figure 2A). Interestingly, whilst *Trib1^mKO^* tumour growth rate accelerated from an early stage and significantly increased from day 18 (p<0.001) and reached 15 mm in diameter (1770 mm^3^ volume) at day 22, *Trib1^mTg^* animals displayed a tumour growth rate similar to wild-type littermates until day 24 (Figure 2B). At this point, tumour volume platoed in wild-type animals, in contrast to *Trib1^mTg^* animals, where tumour continued to grow and reached 15mm in diameter (1770 mm^3^ volume) at day 30 (p<0.01) (Figure 2B). Tumours from both cohorts were collected when they reached 15mm in diameter for further analysis, together with corresponding WT littermate controls.

**Figure 2.**
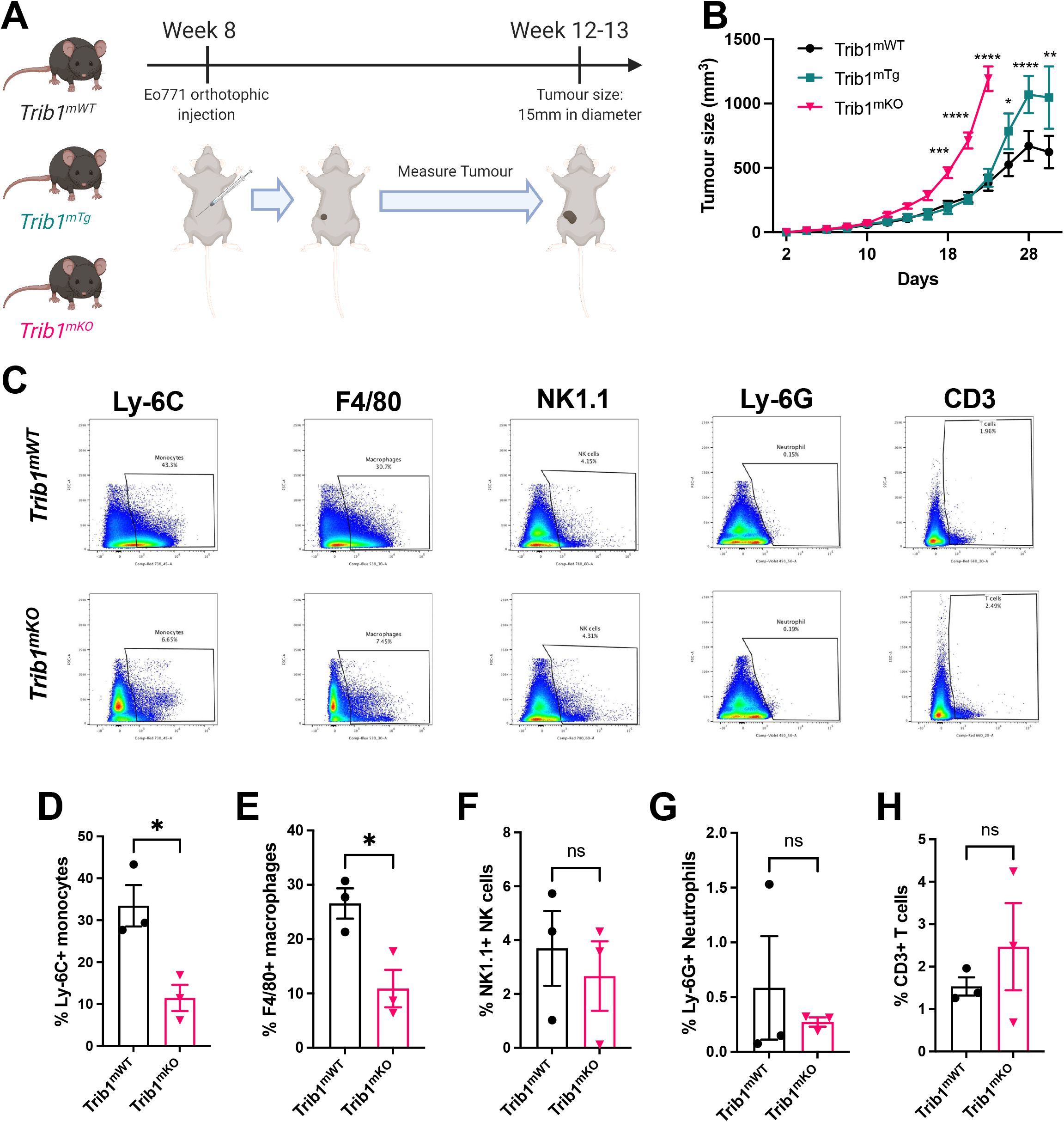
Reduced myeloid *Trib1* expression accelerates breast tumour growth and inhibits myeloid cell infiltration into the TME. (**A**) Development of BC models with altered myeloid *Trib1* expression. Triple-negative, murine BC E0771 cells were injected to the mammary fatpad of myeloid-specific *Trib1* knockout (*Trib1^mKO^*) and transgenic (*Trib1^mTg^*) mice and their respective wild-type (*Trib1^mWT^*) litter-mate controls at the age of 8 weeks. (**B**) Tumour growth in *Trib1^mWT^, Trib1^mTg^* and *Trib1^mKO^* animals was measured and tumour volume calculated. Animals were sacrificed when the tumour reached 15mm in diameter (*Trib1^mKO^* = 22 days; *Trib1^mTg^* = 30 days) and results analysed by two-w ay ANOVA; mean±SEM is plotted; *p<0.05 **p<0.01 ***p<0.001 ****p<0.0001 (n=5-15 mice/group). (**C**) Post-mortem analysis of immune cell content in *Trib1^mKO^* and respective *Trib1^mWT^* tumours by flow cytometry. (**D-H**) Quantification of immune cell content in *Trib1^mKO^* and respective *Trib1^mWT^* tumours. Results of Welch’s t-test are presented; mean±SEM is plotted; *p<0.05 (n=3 mice/group).

### Myeloid-*Trib1* knockout reduces macrophage infiltration and promotes oncogenic cytokine expression in TAMs

Tumours from *Trib1^mKO^* animals were initially analysed by flow cytometry to investigate populations of immune cells in the tumour microenvironment (TME) (Figure 2C, Supplementary figure 2A and C). This analysis revealed that tumours developed in *Trib1^mKO^* animals had a significantly reduced infiltration of both Ly-6C^+^ monocytes and F4/80^+^ macrophages into the TME (p<0.05) (Figure 2D-E). In contrast, the percentage of Ly-6G^+^ neutrophils, NK1.1^+^ NK cells, and CD3^+^ T-cellsand its subtypes (CD4^+^ naïve and CD8^+^ cytotoxic T cells) in the tumour were not altered between *Trib1^mWT^* and *Trib1^mKO^* (Figure 2F-H, Supplementary figure 2B and D).

In order to explore the potential mechanisms of the observed accelerated mammary tumour growth and its links with the reduced monocyte and macrophage infiltration in *Trib1^mKO^* mice, we further assessed the localisation of macrophages and their phenotype using fluorescence staining and flow cytometry (Figure 3A and 3C, and Supplementary figure 2E). Perivascular TAMs (PV TAMs) are in close contact with blood vessels (within 250μm radius) and play a crucial role in angiogenesis of mammary cancers as well as metastasis and intravasation of cancer cells (Hughes et al., 2015, Arwert et al., 2018). Staining of TAMs and endothelial cells with F4/80 and CD31, respectively, revealed a significant reduction of PV TAMs in *Trib1^mKO^* tumours (Figure 3B). However, although an increase in pro-inflammatory macrophages has been reported previously both in full-body and myeloid-specific *Trib1* knockout animals (Satoh et al., 2013, Johnston et al., 2019), inhibition of myeloid *Trib1* expression did not alter the ratio of NOS2^+^ pro-inflammatory TAMs and mannose receptor (MR)^+^, anti-inflammatory TAMs in the TME (Figure 3D, Supplementary figure 2F), suggesting that the reduced TAM numbers rather their phenotype contributed to the accelerated tumour growth.

**Figure 3.**
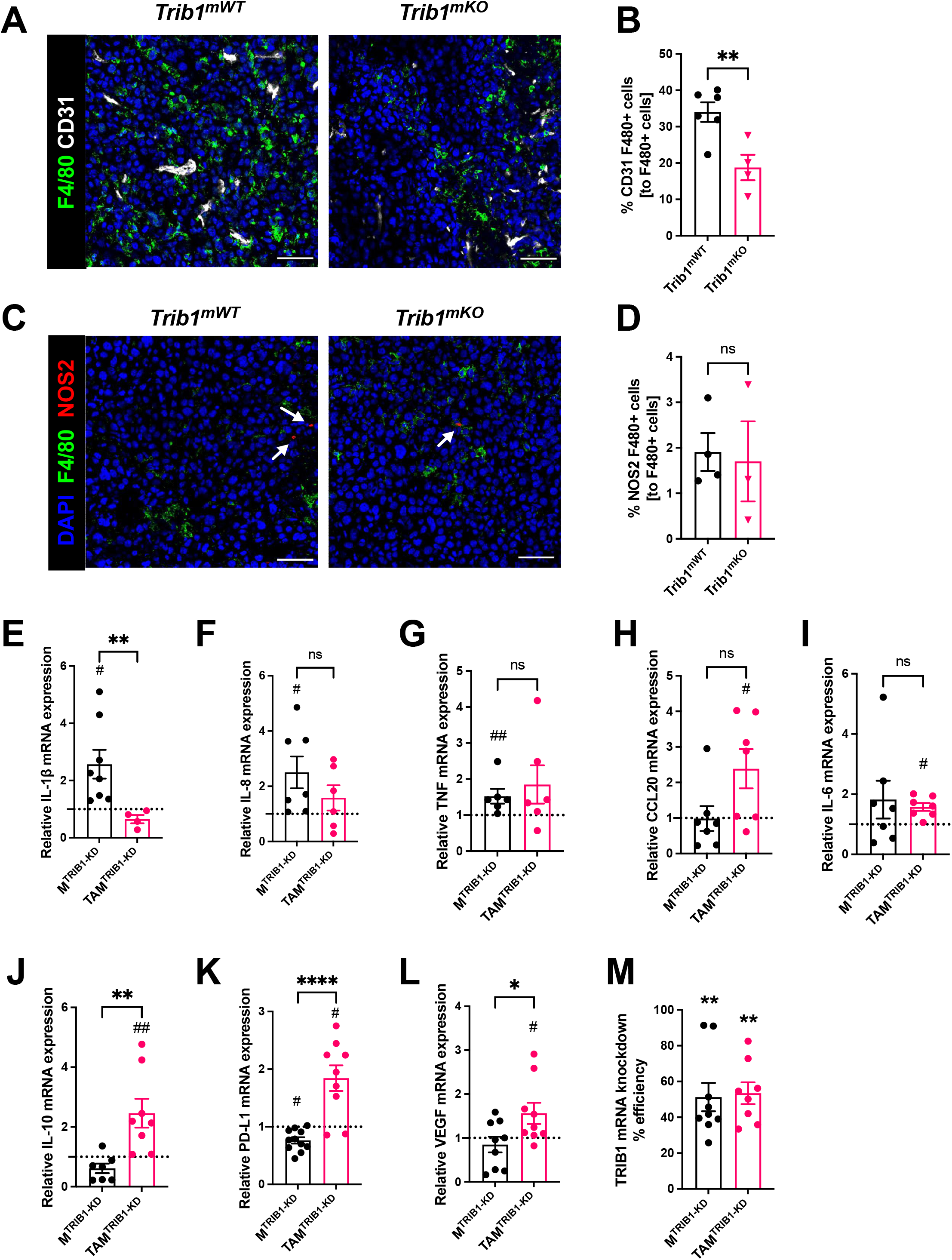
Myeloid *Trib1* knockout inhibits perivascular TAM infiltration, and *TRIB1* knockdown in TAMs enhances oncogenic cytokine expression. (**A**) Representative images of CD31 (white) and F4/80 (green) fluorescence staining in *Trib1^mKO^* and respective *Trib1^mWT^* tumours (Scale: 50μm); **(B)** Quantification of perivascular TAMs (F4/80+ CD31+) in *Trib1^mKO^* tumours relative to the total number of F4/80+ TAMs. Cell numbers were quantified manually from 5 randomly taken field of views using ImageJ. Results of unpaired t-test are presented; mean±SEM is plotted; **p<0.01 (n=4-6 mice/group); **(C)** NOS2 (red) and F4/80 (green) fluorescence staining in *Trib1^mKO^* and respective *Trib1^mWT^* tumours (Scale: 50μm). NOS2 (red) staining is marked with white arrows.; **(D)** Quantification of pro-inflammatory TAMs (F4/80+ NOS2+) in *Trib1^mKO^* tumours relative to the total number of F4/80+ TAMs. Cell numbers were quantified manually from 5 randomly taken field of views using ImageJ. Results of unpaired t-test are presented; Mean±SEM is plotted (n=3-4mice/group). **(E-M)** Human MDMs isolated and differentiated from blood were transfected with either non-targeting control or *TRIB1* siRNA for 48 hours and either left unpolarised or polarised to TAMs using the CM for 24 hours. Expression values were normalised to non-targeting control and expressed as a fold difference to this (dotted line). RNA levels of *IL-1β, IL-8, TNF, CCL20, IL-6, IL-10, PD-L1*, and *VEGF* (**E-L**) and efficiency of *TRIB1* siRNA transfection (**M**) 48 hours after *TRIB1* siRNA transfection (M^TRIB1-KD^) and TAM polarisation (TAM^TRIB1-KD^) were assessed. Results of paired and unpaired t-tests are presented; mean±SEM is plotted; *p<0.05 **p<0.01 ****p<0.0001 represent p-values between M^TRIB1-KD^ *vs*. TAM^TRIB1-KD^ whilst #p<0.05 ##p<0.005 represent p-values of M^TRIB1-KD^ and TAM^TRIB1-KD^ to their respective control (M^Control^ and TAM^Control^) (n=4-9 donor/group).

In order to gain a mechanistic insight into how *TRIB1* regulates monocyte-derived macrophages and its impact on re-educating macrophages towards TAMs, human MDMs were transfected with siRNA to reduce *TRIB1* levels (M^TRIB1-KD^), followed by a treatment with tumour-conditioned medium (CM) (TAM^TRIB1-KD^). Expression of key cytokines were assessed by RT-qPCR, revealing that expression of pro-inflammatory cytokines *IL-1β* (p<0.05), *IL-8* (p<0.05), and *TNF* (p<0.01) were significantly increased in M^TRIB1-KD^ but were not altered in TAM^TRIB1-KD^, compared to non-targeting siRNA transfected MDMs (Figure 3E-G, Supplementary figure 4A-C and I-K). In contrast, *TRIB1* knockdown in TAM^TRIB1-KD^ significantly induced expression of several oncogenic cytokines, including *CCL20* (p<0.05), *IL-6* (p<0.05), *IL-10* (p<0.01), *PD-L1* (p<0.05), and *VEGF* (p<0.05), compared to M^TRIB1-KD^ (Figure 3E, Supplementary figure 4D-H and L-P). Notably, *IL-10* (p<0.01), *PD-L1* (p<0.0001) and *VEGF* (p<0.05) expression were significantly increased in TAM^TRIB1-KD^ compared to M^TRIB1-KD^ (Figure 3E), suggesting the myeloid *TRIB1* is an important regulator of oncogenic cytokine expression in TAMs, downstream of signals secreted by tumour cells.

### Overexpression of *Trib1* reduces hypoxic TAM infiltration and inhibits pro-inflammatory TAM polarisation

Analysis of tumour growth in *Trib1^mTg^* mice (Figure 2A) revealed that elevated myeloid-*Trib1* levels lead to an increase in tumour size at advanced stages. To gain a mechanistic understanding of this effect, TAM localisation and phenotypes in *Trib1^mTg^* TME were investigated using fluorescence staining. Carbonic anhydrase IX (CA9) is a cell-surface glycoprotein in the tumour, expression of which is induced by hypoxia and has been shown to be involved in cancer progression (Pastorekova and Gillies, 2019). Thus, staining for CD31 and CA9 were used together with F4/80 to identify PV TAMs (Figure 4A-C, and Supplementary figure 5) *vs*. TAMs residing in hypoxic areas (Figure 4D, E). Pro- and antiinflammatory markers, NOS2 and MR, were used to characterise TAM phenotypes in *Trib1^mTg^* tumours (Figure 4F-K). Similar to *Trib1^mKO^*, we observed a significant overall reduction of F4/80+ TAM numbers in *Trib1^mTg^* (p<0.05) (Figure 4B). Whilst there was no difference in CD31^+^ F4/80^+^ PV TAMs (Figure 4C), a significant reduction was observed in CA9^+^ F4/80^+^ hypoxic TAMs in *Trib1^mTg^* tumours, compared to *Trib1^mWT^* (p<0.005) (Figure 4E). Further, staining of TAMs with phenotypic markers (NOS2 and MR) also demonstrated a reduction in NOS2^+^ TAM numbers (p<0.005), including PV TAMs (p<0.05) (Figure 4F-H) but did not alter the ratio of MR^+^ TAMs (Figure 4I-K).

**Figure 4.**
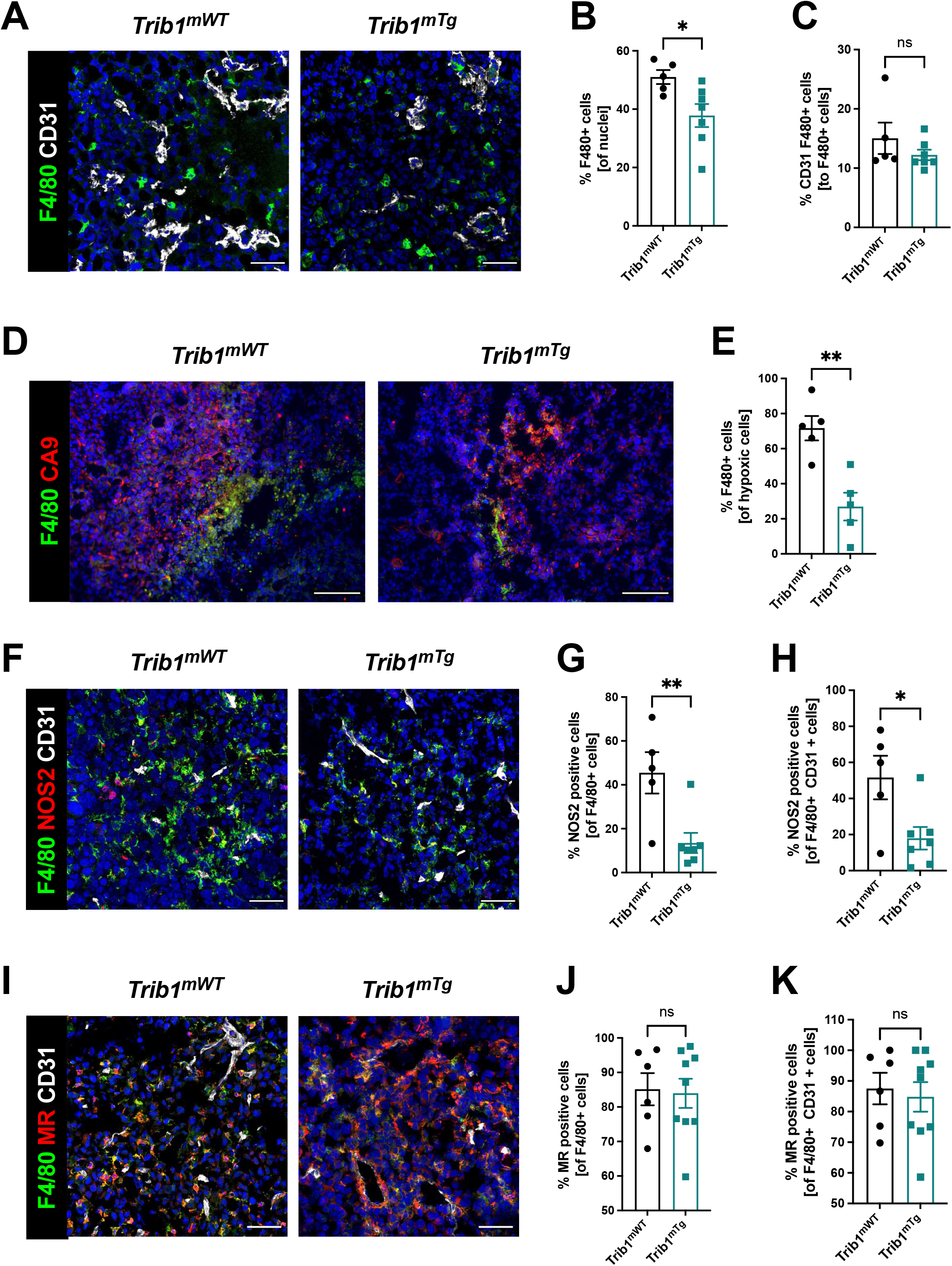
Overexpression of myeloid *Trib1* reduces hypoxic TAM numbers in the TME and inhibits TAM polarisation towards a pro-inflammatory phenotype. **(A)** Representative images of CD31 (white) and F4/80 (green) fluorescence staining (Scale: 50μm). Cells were quantified manually from 4-5 randomly taken fields of view using ImageJ. **(B)** Quantification of TAMs and **(C)** perivascular TAMs in *Trib1^mTg^* and respective *Trib1^mWT^* tumours. **(D)** Representative images of CA9 (red) and F4/80 (green) fluorescence staining on *Trib1^mWT^* and *Trib1^mTg^* tumours (Scale: 100μm). Cells were quantified manually from 4-5 randomly taken fields of view using ImageJ. **(E)** Quantification of TAMs in the hypoxic area relative to total cell numbers in hypoxia. **(F)** Representative images of CD31 (white), NOS2 (red), and F4/80 (green) fluorescence staining on tumours (Scale: 50μm). Cells were quantified manually from 4-5 randomly taken field of views using ImageJ. **(G)** Quantification of pro-inflammatory TAMs and **(H)** pro-inflammatory perivascular TAMs in *Trib1^mTg^* and respective *Trib1^mWT^* tumours. **(I)** Representative images of CD31 (white), MR (red), and F4/80 (green) fluorescence staining on tumours (Scale: 50μm). Cells were quantified manually from 4-5 randomly taken field of views using ImageJ. **(J)** Quantification of anti-inflammatory TAMs and (**K**) anti-inflammatory perivascular TAMs in *Trib1^mTg^* and respective *Trib1^mWT^* tumours. Results of unpaired t-test are presented; mean±SEM is plotted; *p<0.05 **p<0.01 (n=5-9 mice/group).

### *Trib1^mTg^* tumours display reduced T cell infiltration and reduced IL-15 expression

Recruitment of T-cells to the TME is a central mechanism for inhibition of tumourigenesis and cytokines secreted by TAMs play a key role in this process (Bogen et al., 2019, Jeong et al., 2019). Our above data demonstrates that reduced *vs*. elevated expression of *mTrib1* leads to distinct changes in TAM phenotypes and have also shown that recruitment of T-cells in *Trib1^mKO^* tumours is unaltered (Figure 2H and Supplementary Figure 2A-D). Thus, we next tested whether the observed alterations in TAM numbers and phenotypes affect T-cell recruitment in *Trib1^mTg^* animals. Fluorescence staining was used to identify changes in CD3^+^ T-cell numbers and populations of CD4^+^ *naïve* and CD8^+^ cytotoxic T-cells (Figure 5A and B, Supplementary figure 6), revealing a significant reduction in the overall number of CD3^+^ T-cells in the *Trib1^mTg^* TME, compared to *Trib1^WT^* (p<0.01) (Figure 5C). Furthermore, the proportion of both CD4^+^ CD3^+^ *naïve* T-cells and CD8^+^ CD3^+^ cytotoxic T-cells was significantly reduced in *Trib1^mTg^* (p<0.05) (Figure 5D-E).

**Figure 5.**
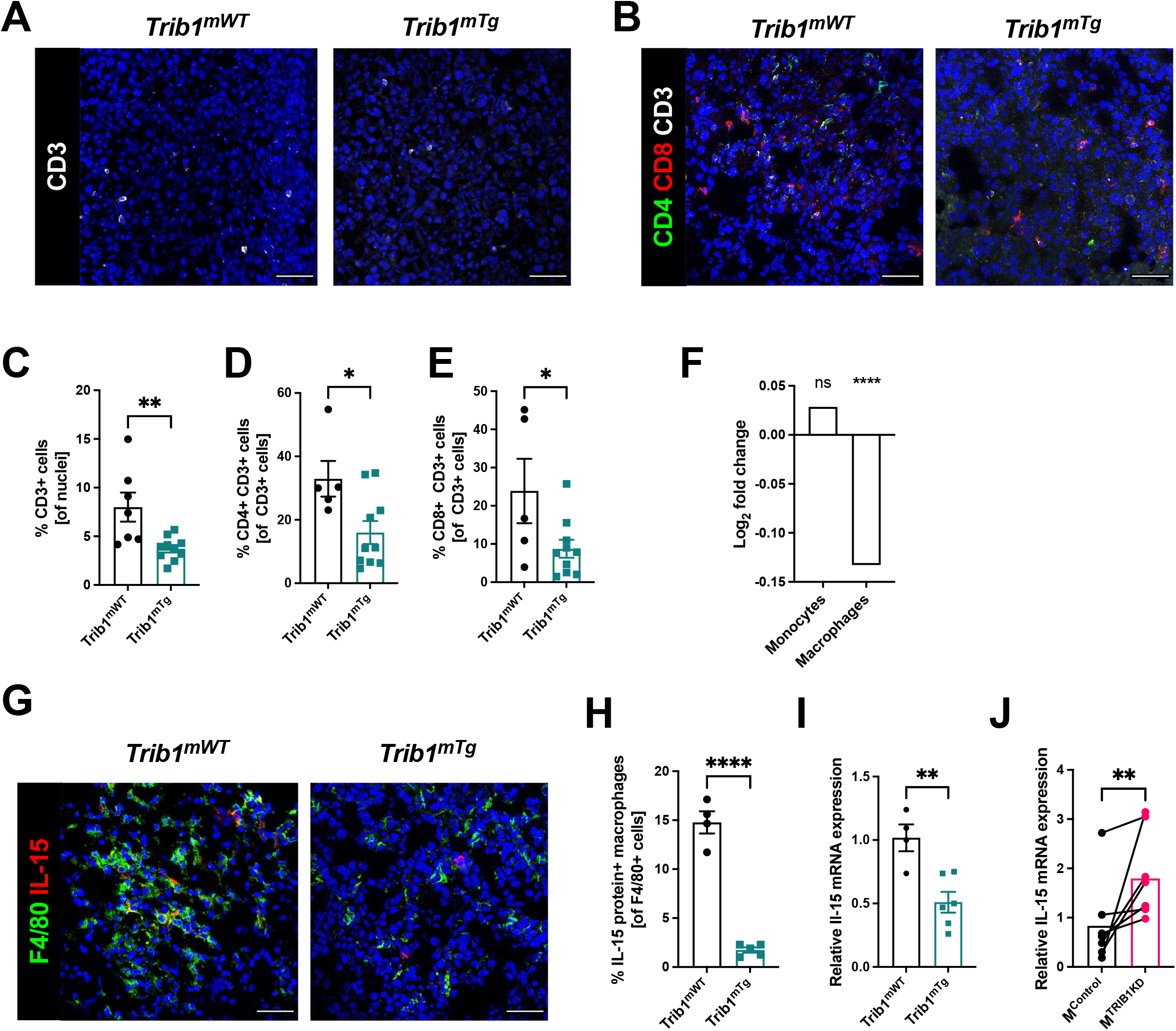
Myeloid *Trib1* overexpression impairs IL-15 expression and significantly reduces T-cellsin the TME. **(A–B)** Representative images of CD3 (white) staining in *Trib1^mTg^* and respective *Trib1^mWT^* tumours (Scale: 50μm); and CD3 (white), CD4 (green), and CD8 (red) fluorescence staining in *Trib1^mTg^* and respective *Trib1^mWT^* tumours (Scale: 50μm). Cells were quantified manually from 4-5 randomly taken field of views using ImageJ. **(C)** Quantification of T-cellsin *Trib1^mTg^* and respective *Trib1^mWT^* tumours. Results of unpaired t-test is presented; mean±SEM is plotted; **p<0.01 (n=7-10 mice/group). **(D-E)** Quantification of CD4+ *naïve* and CD8+ cytotoxic T-cellsin *Trib1^mTg^* and respective *Trib1^mWT^* tumours. Results of unpaired t-test are presented; mean±SEM is plotted; *p<0.05 (n=5-11 mice/group). **(F)** *IL-15* expression in human monocytes and MDMs from participants of the Cardiogenics Transcriptomic Study. MDM (n=596) and monocytes (n = 758) were ranked according to *TRIB1* RNA levels and *IL-15* expressed between the top vs. bottom 25% of the samples were plotted. Results of FDR adjusted p-values are presented as ****p<0.0001. **(G)** Representative images of IL-15 (red) and F4/80 (green) fluorescence staining in *Trib1^mTg^* and respective *Trib1^mWT^* tumours (Scale: 50μm). Cells were quantified manually from 4 randomly taken field of views using ImageJ. **(H)** Quantification of TAMs expressing IL-15 in *Trib1^mTg^* and respective *Trib1^mWT^* TME relative to the total number of TAMs. Results of unpaired t-test is presented; mean±SEM is plotted; ****p<0.0001 (n=4-5 mice/group). **(I)** Mouse BMDMs isolated from *Trib1^mWT^* and *Trib1^mTg^* animals were analysed with RT-qPCR. RNA level of *Il-15* in mouse BMDMs. Results of unpaired t-test is presented; mean±SEM is plotted; **p<0.01 (n=4-6mice/group). **(J)** Human MDMs isolated and differentiate from blood were transfected with either control or *TRIB1* siRNA for 48 hours and *IL-15* RNA expression was analysed. Results of paired t-test is presented; mean is plotted; **p<0.01 (n=6 donor/group).

In order to identify *mTrib1*-dependent mechanisms that may explain an impaired T-cell recruitment to the tumour, we have assessed the correlation between *TRIB1* levels and genes that have been shown to regulate T-cell recruitment in the Cardiogenics Transcriptomic Study (Heinig et al., 2010, Schunkert et al., 2011, Rotival et al., 2011). This analysis revealed that high *TRIB1* levels correlate very significantly with a reduced IL-15 expression in human macrophages (but not in monocytes) (Figure 5F). IL-15 is a cytokine expressed by myeloid cells crucial for the development, function and survival of T-cells. IL-15 stimulates tumour-specific T-cell responses, increases cellular growth, inhibits apoptosis, and enhances immune cell activation, and as a consequence, promotes anti-tumour responses (Robinson and Schluns, 2017). Thus, we next assessed IL-15 in *Trib1^mTg^* TAMs by fluorescence staining (Figure 5G, Supplementary figure 6). Quantification of IL-15^+^ TAMs revealed that overexpression of *Trib1* led to significantly reduced IL-15 expressing TAM numbers in the TME (p<0.01) (Figure 5H). To substantiate that *TRIB1* is a direct regulator of *IL-15* expression in myeloid cells, RT-qPCR analysis was performed in BMDMs isolated from *Trib1^mTg^* showing that enhanced *Trib1* expression led to a significant decrease in *IL-15* expression (Figure 5I). In contrast, transient knockdown of *TRIB1* with siRNA transfection in human MDMs from healthy human participants significantly increased *IL-15* expression (p<0.01) (Figure 5J).

## Discussion

It is now widely recognised that TAMs are a crucial component of TME and the number of these cells is associated with cancer cell resistance to therapy, poor patient survival and prognosis (Zhang et al., 2013, Yuan et al., 2014, Zhao et al., 2017, Qiu et al., 2018). However, the molecular mechanisms that shape TAM phenotype and thus determine whether they are pro-tumorigenic or promoting anti-tumour immune responses, are poorly understood.

A pseudokinase protein, TRIB1, has been reported as a potential regulator of macrophage phenotypes. It is highly expressed in the myeloid lineage and is associated with altered tissue macrophage phenotypes (Satoh et al., 2013, Johnston et al., 2019, Richmond and Keeshan, 2020). Although the effect of *TRIB1* in TAMs has not been elucidated, previous studies investigated *TRIB1* as an oncogene in prostate and colon cancer and also associated it with sensitivity of breast cancer cells to Tumor necrosis factor-related apoptosis-inducing ligand (TRAIL) -induced apoptosis (Gendelman et al., 2017, Wang et al., 2017, Shahrouzi et al., 2020). Putting together these published data, our observations that *TRIB1* expression is associated with patient survival and therapy responses in breast cancer patients and that the majority of TAMs highly express TRIB1 in a murine model of BC, we hypothesised that TRIB1 expression in myeloid cells may alter TAM phenotypes. As a consequence, *mTRIB1* would mechanistically contributes to breast cancer tumourigenesis, as well as to response to chemotherapy.

Of note, the prediction of response is only observed in those tumours with a higher rate of proliferation, such as the basal-like and luminal B phenotypes. The latter is characterised by the dual expression of the estrogen and HER2 receptor what constitutes two druggable oncogenic vulnerabilities (Baliu-Pique et al., 2020). Chemotherapy, and particularly anthracyclines, is a backbone treatment in this disease and its known that the immunologic state can modulate the efficacy to these agents through the presence of different immune populations and secreted factors (Perez-Pena et al., 2019).

We used E0771 cells in this study, that have been shown to express estrogen receptor (ER), progesterone receptor (PR) and ERBB2 and classified into luminal B subtype, which is found in 30-40% of BCs and generally known to be more aggressive than luminal A BCs (Le Naour et al., 2020). Of note, our analysis of patient survival showed that *TRIB1* expression is elevated in tumours responding to chemotherapy in Luminal B BC, compared to non-responders, further justifying the choice of this murine model.

TAM phenotype in the solid tumour is critical for tumour growth, where proteins secreted by cancer cells (such as IL-4, IL-10, and CSF-1) drive TAMs towards an anti-inflammatory phenotype that promotes angiogenesis and immunosuppression (Yuan et al., 2015, Chen et al., 2019b). However, pro-inflammatory TAMs can also play oncogenic roles, particularly at the early phases of tumour growth, linked to hypoxia. Abundant infiltration of pro-inflammatory TAMs has been observed in early tumour development, and the expression of TNF and activation of PGC-1α and AMPK was shown to promote glycolysis and exacerbate tumour hypoxia (Pinto et al., 2019, Jeong et al., 2019). The importance of hypoxic signals has also been evidenced where knockout of HIF-1α reduced the proliferation of BC cells *in vitro* as well as primary breast tumour volume by 60% *in vivo* (Schwab et al., 2012). Hypoxic TAMs have also been shown to secrete angiogenic proteins, with HIF-1α stimulating pro-angiogenic functions in TAMs, thus facilitating tumour vascularisation (Henze and Mazzone, 2016, Chen et al., 2019b). Perivascular (PV) TAMs express high levels of MRC1 and VEGF to facilitate tumour angiogenesis, and help formation of paracrine feedback loops (CSF1 from cancer cells, EGF from TAMs, and HGF from endothelial cells) to initiate metastasis and intravasation of cancer cells at the TME of metastasis (Claire et al., 2016, Hughes et al., 2015, Arwert et al., 2018, Lapenna et al., 2018). Thereby, although *Trib1^mKO^* and *Trib1^mTg^* both demonstrated a significant reduction in TAM infiltration overall in our BC models, analysis of *Trib1^mTg^* revealed a significant and localised reduction of TAM numbers in hypoxic areas, as well as inhibition of pro-inflammatory TAM polarisation in the TME, both of which mechanisms that may contribute to the observed late acceleration of mammary tumour growth.

In contrast, *Trib1^mKO^* reduced infiltration of PV TAMs but did not alter the number of NOS2 positive macrophages in the tumour. Instead, *in vitro TRIB1* knockdown in a model of human TAMs revealed that inhibition of *TRIB1* enhances expression of oncogenic cytokines in TAMs, which are involved both in cancer cell survival and immune suppression. Increased IL-6 expression in TAMs was reported to promote cancer cell survival resistance to hypoxia (Jeong et al., 2017); IL-10 is known to suppress immune surveillance, inhibit apoptosis, and to enhance migration of cancer cells (Sheikhpour et al., 2018, Chen et al., 2019a); overexpression of PD-L1 disrupts T-cell proliferation and function (Wu et al., 2019) and VEGF enhances BC growth and angiogenesis (Riabov et al., 2014). These observations are in line with our previous work, where we have shown that myeloid *Trib1*-deficiency alters macrophage function (in that case, formation of foam cells in the atherosclerotic plaque), rather than a clear shift in inflammatory status of *Trib1^mKO^* cells (Johnston et al., 2019).

TAMs interact with T-cellsin TME to suppress T cell-driven cytotoxic immune response and promote tumour growth. Previous studies reported that TAMs impair CD8^+^ T-cell activation and proliferation, and depletion of TAMs in the TME enhances the infiltration of both *naïve* and cytotoxic T-cells (Williams et al., 2016, Jeong et al., 2019). Tumour-infiltrating T-cells enter tumour at an early stage as *naïve* CD4^+^ T-cells, followed by macrophage infiltration and contribute to early tumour rejection and/or anti-tumour effects through promoting senescence and tumour apoptosis via secretion of cytokines (such as IFNγ and TNF) and interact with macrophages, NK cells and CD8^+^ T-cells to enhance tumour eradication (Bogen et al., 2019). Interstingly, a recent study from Carrero *et al*. have shown that most myeloid cells in the tumour TME express high levels of IL-15 (Santana Carrero et al., 2019), proposing that these stromal cells may be a critical source of this anti-tumour cytokine. In this study, we identified a significantly reduced infiltration of both CD4^+^ *naïve* and CD8^+^ cytotoxic T-cellsinto the TME in *Trib1^mTg^* animals, despite TAM infiltration being inhibited. Our mechanistic analysis revealed that regulation of T cell infiltration may be due to a previously unrecognised role of myeloid-TRIB1 as a critical regulator of IL-15 expression.

Given the data we present in this study, we propose that dysregulated levels of *TRIB1* in myeloid cells lead to accelerated tumour growth via distinct molecular mechanisms (Figure 6). More generally, this study exemplifies how alterations in the expression of the same gene in TAMs may have opposing consequences at different stages of tumour development. Whilst it is to be formally tested in future studies, we speculate that *TRIB1* expression changes in TAMs could be associated with the initiaton and/or with the growth of the tumor and adaptation to lack of nutrients, as well as to hypoxic environment. Nevertheless, knockout of myeloid-*TRIB1* upregulates the expression of oncogenic cytokines in TAMs whilst its overexpression modifies TAM phenotype and T-cell composition in the TME, both enhancing tumour growth. Such data reinforce the general concept for the complex role of TAMs in BC and analysis of consequences for altered *TRIB1* expression highlight potential diagnostic/prognostic markers and therapeutic markers for anti-cancer immunotherapy. In addition, our findings also support the idea that enhanced *TRIB1* expression could be explored as a potential biomarker in BC that might help to predict response chemotherapy.

**Figure 6.**
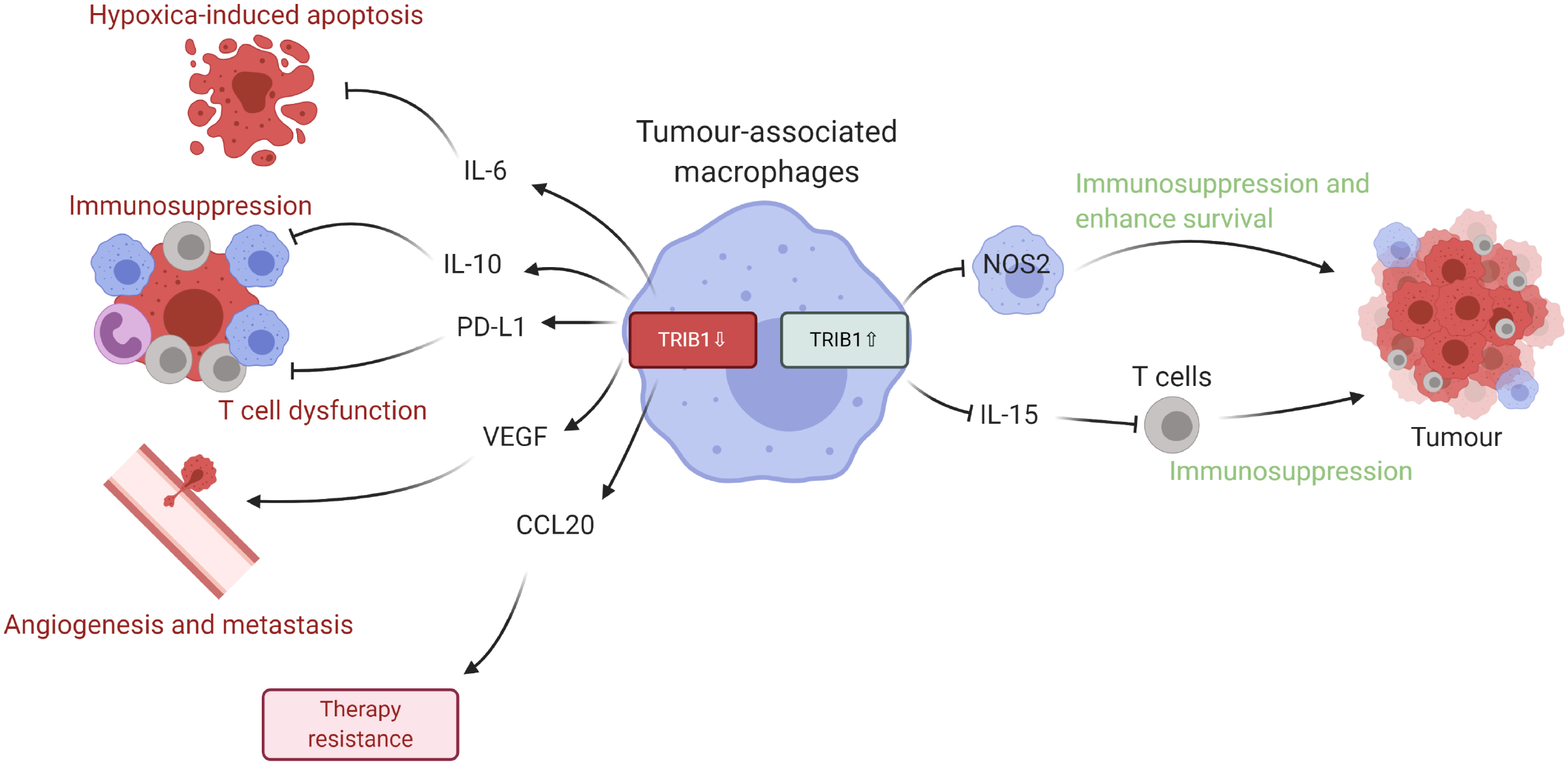
Schematic representation of TRIB1 mediated TAM regulation. TRIB1 distinctly regulate TAMs to enhance tumour growth. Reduction of TRIB1 accelerates oncogenic cytokine IL-6, IL-10, CCL20, PD-L1, and VEGF expressions in TAMs to inhibit apoptosis and enhance immune suppression, angiogenesis, and therapeutic resistance. Whilst the overexpression inhibits pro-inflammatory phenotype TAM in the TME to suppress the immune response and enhance cancer cell survival and decrease IL-15 expression, which then reduces T cell infiltration and disrupts T cell-induced immune responses. The image was created with BioRender.com.

### STAR Methods

#### Study approval

All animal studies were approved and conducted in accordance with the University of Sheffield code of ethics and Home Office regulations (project license No.PPL70/8670). Human monocyte-derived macrophages (MDMs) isolated from healthy participants were obtained with signed participant-informed consent and approval from the University of Sheffield Research Ethics Committee (project license No.SMBRER310) and in accordance with the Declaration of Helsinki. All participants gave written informed consent..

#### Microarray analysis

Cardiogenic Transcriptomic Study (Schunkert et al., 2011, Heinig et al., 2010, Rotival et al., 2011) was analysed in collaboration with Professor Alison Goodall and Dr Stephen Hamby at the University of Leicester. The analysis was performed as described in (Johnston et al., 2019). In brief, top and bottom quartiles of TRIB1 expressing monocytes (N = 758) and macrophages (N = 596) were compared and obtained the TRIB1 co-regulated, differentially expressed genes using FDR adjusted p-values of <0.01, cut-off log-2 fold changes of >0.071 (upregulated) and >-0.071 (down-regulated). The gene list was further analysed with QuSage (Yaari et al., 2013, Turner et al., 2015, Meng et al., 2019) to identify the pathways enriched in the TRIB1 coexpressed gene sets.

#### Mice

All mice were bred on a *C57BL/6* genetic background under the University of Sheffield code of ethics, and Home Office regulations in the University of Sheffield Biological Service Unit. Trib1 fl/fl x Lyz2Cre (*Trib1^mKO^*), ROSA26.Trib1Tg x Lyz2Cre (*Trib1^mTg^*) and their corresponding WT controls have recently been described (Johnston et al., 2019).

#### Tumour models

The mouse E0771 cell line (obtained from Dr Jessalyin Ubellacker (University of Harvard, USA) was cultured in DMEM medium (Gibco) containing 10% (v/v) low endotoxin heat-inactivated foetal bovine serum (Biowest), and 1% L-glutamine (Lonza). Eight-week-old female *Trib1^mKO^* and *Trib1^mTg^* mice were inoculated with 3 x 105 E0771 cells into the right nipple via intra-ductal injection. Once the tumours formed, the size was measured every 2 days with callipers until it reached 15 mm in diameter.

#### Cancer cell culture and conditioned medium production

MDA-MB-231, BT474, SKBR3 and MCF7 cell lines were cultured in RPMI-1640 (Gibco) with 10% (v/v) low endotoxin heat-inactivated foetal bovine serum (LE-FBS)(Biowest), 1% (v/v) streptomycin/penicillin (Gibco), 1% L-glutamine (Lonza). All cells were obtained from Dr. Penelope Ottowell and Dr. Munitta Muthana (University of Sheffield, UK) and subsequently maintained in our laboratory. They were routinely tested for mycoplasma contamination and all tested negative. To obtain MDA-MB-231 conditioned medium (CM), cells were cultured in T75 flasks for 48 hours at 37C in a 5% CO2 and centrifuged at 600 x g for 5 minutes to remove cells and cellular debris.

#### Isolation of human blood monocytes

Whole blood was collected in 3.8% trisodium citrate (Sigma) and used immediately to isolate cells. In 15 ml of Ficoll-Paque PLUS (GE Healthcare), 30 ml of blood was gently layered and centrifuged at 900 x g for 20 minutes at room temperature (RT) to separate peripheral blood mononuclear cells (PBMCs) from plasma. PBMCs were recruited in PBS–EDTA (Thermo Fischer) solution (PBSE) and centrifuged at 400 x g for 5 minutes at RT. After red blood cell lysis with 10 ml of RBC lysis buffer (155 mM NH4Cl, 10mM KHCO3, 0.1M EDTA in H2O) for 5 minutes at RT, 40 ml of PBSE was added and centrifuged at 1500 rpm or 400 x g for 5 minutes. Cells were counted using a haemocytometer (Hawksley) and resuspended in 90 μl 4°C MACS buffer (0.5% [w/v] bovine serum albumin [BSA, Sigma] – PBSE) and 10 μl CD14+ microbeads (Miltenyi Biotec) per 10^7 cells for 15 minutes at 4°C. 2 ml of MACS buffer was added and centrifuged at 260 x g for 5 minutes. CD14+ monocytes were isolated with LS column (Miltenyi Biotec) and MidiMACSTM Separator (Miltenyi Biotec) for differentiation.

#### TRIB1 siRNA transfection

Viromer Green (Lipocalyx) was used to transfect TRIB1 siRNA (ON-TARGET plus siRNA, Dharmacon) and Non-Targeting Control siRNA (ON-TARGET plus siRNA, Dharmacon) in order to knockdown TRIB1 level in humans MDMs according to the manufacturer’s instructions.

#### Monocyte-derived macrophages differentiation and stimulation

Isolated monocytes were incubated in fresh medium (RPMI-1640 (Gibco) 10% (v/v) LE-FBS (Biowest), 1% (v/v) streptomycin/penicillin (Gibco), 1% L-glutamine (Lonza)) with 100 ng/ml recombinant human (rh) macrophage-colony stimulating factor (M-CSF) (Peprotech) for 7 days at 37°C at 5% CO-2 to facilitate differentiation of monocytes to macrophages. MDMs were washed with PBS and polarised by incubating with 20 ng/ml IFN-γ (Peprotech) and 100 ng/ml E. coli lipopolysaccharide (Serotype R515 TLRgradeTM, Enzo Life Sciences), 20 ng/ml IL-4 (Peprotech), 20 ng/ml IL-10 (Peprotech), and CM for 24 hours at 37°C at 5% CO-2.

#### Isolation of BMDMs

The femur and tibias of mice were collected, and tissues were gently removed from the bones. Bone marrow was harvested by flushing the bones with PBS using a 2.5 ml syringe. Any clumps of cells was dispersed with a pipette and passed through 70μm cell strainer (Fisher Scientific). The cell suspension was centrifuged at 500 x g for 5 minutes, and the pellet was cultured in fresh L929 cell-conditioned DMEM medium for 6 days to differentiate into BMDMs.

#### Protein extraction and quantification

Cells washed with PBS were collected into a 1.5ml Eppendorf tube and thoroughly mixed with lysis buffer (RIPA buffer with 1% protease and phosphatase inhibitor). Cells were incubated at −80°C for 30 minutes and sonicated for 15 seconds to allow further lysis. Cells were then centrifuged at 15,000 x g for 10 minutes at 4°C to remove debris and supernatant was collected and stored at −80°C. PierceTM BCA Protein Assay Kit (Thermo Scientific) was used to quantify proteins as manufacturer’s instructions.

#### Western blot

Proteins were mixed with 5x laemmli buffer and incubated at 100°C for 10 minutes. Samples were immediately transferred in the ice afterwards. All samples and prestained protein ladder (10-250 kDa, Thermo ScientificTM) were loaded into the columns of NuPAGETM 4-12% Bis-Tris Gel (Invitrogen) placed in the Invitrogen tank containing 1x NuPAGE MOPS SDS running buffer (Novex). The gel was run at 100v for 75 minutes and transferred to a PVDF (Polyvinylidene difluoride) membrane (Millipore) using NuPAGE transfer buffer (Novex) with methanol and antioxidant (Invitrogen) at 35v for 60 minutes. The membrane was blocked with 5% milk-TBST at RT for 1 hour and incubated overnight with TRIB1 (Millipore), and HSP90 (Abcam) diluted in 5% milk-TBST (1:1000 and 1:5000 respectively) at 4°C. The membrane was then washed with 0.1 v/v TBST for 5 minutes 3 times and incubated with Polyclonal Goat anti-Rabbit Immunoglobulin/HRP, and Polyclonal Rabbit anti-Rat Immunoglobulin/HRP (Dako) diluted in 5% milk-TBST (1:2500 and 1:5000 respectively) at RT for 1 hour. The membrane was then washed with TBST 3 times for 5 minutes, incubated with ECL, and imaged with Bio-Rad imager.

#### RNA extraction and quantification

Cells were gently washed twice in PBS and incubated at RT for 5 minutes in 700 μl of QIAzol lysis reagent to homogenate the cells, 140μl of chloroform was added to cells and RNA extracted using the miRNeasy Mini Kit (Qiagen) according to the manufacturer’s instructions. The amount of RNA was quantified using a Nanodrop Spectrophotometer ND1000 and used for RT-qPCR analysis or stored at −80°C.

#### cDNA synthesis and Real-time quantitative PCR analysis

cDNA was produced with iScript cDNA synthesis kit (Bio-Rad) according to the manufacturer’s instructions. Quantitative RT-PCR was performed using primers designed with NCBI BLAST to target human macrophage polarisation markers (Supplementary table 1) and PrecisionPLUS SYBR-Green master mix (Primerdesign). 364-well plate was used in the experiment to assess the expression of genes in the samples. SYBR green master mix with forward and reverse primers were added to each well of the 364 well RT-qPCR plates at a total volume of 5.6μl. Followed by the addition of 5μl of cDNA (0.4 ng/ul) and the plate was centrifuged for 2 minutes at 2000rpm. The RT-qPCR plate was then assessed on a Bio-Rad I-Cycler PCR machine with the protocol provided by the manufacturer. GAPDH and B-actin were used as the housekeeper, and the changes in gene expression were obtained using the 2^-ΔΔCT^ method.

#### Tissue dissociation

The tumour tissue collected from mice was shredded with scissors, and placed in 5 ml of tumour-dissociation medium (TDM) (IMDM medium, 0.2 mg/ml collagenase IV, 2 mg/ml dispase, 1.25 ug/ml DNase 1) in a 15ml bijou tube, and rotated at 37°C for 30 minutes. 5 ml of 10% FBS-TDM was added into the tube and passed through a 70μm filter (Fisher Scientific). The samples were placed directly on ice and centrifuged at 4500 rpm for 5 minutes. The cell pellet was washed three times with PBS and used for flow cytometry analysis.

#### Flow cytometry

Total tumour cells were resuspended in PBS and centrifuged at 500 x g for 5 minutes. The samples were resuspended in 100μl LIVE/DEAD Fixable Blue Dead Cell Stain kit (Invitrogen) and incubated for 15 minutes at RT in the dark, and 200μl of PBS was added to the tube and centrifuged at 500 x g for 5 minutes. The pellet was aliquot with PBS if required and centrifuged at 500 x g for 5 minutes. Cells were stained with following antibodies at 1:25 dilution: F4/80 Alexa Fluor 488 (Bio-rad); 1:100 dilutions: CD3 APC (Tonbo Bioscience), MR PE, Ly-6C Alexa Fluor 700, NK1.1 APC-Cy7, Ly-6G Pacific Blue, CD4 PerCP/Cy5.5, CD8 APC-Cy7, CD279 PE (Biolegend); 1:200 dilution: CD274 PE-Cy7 (Biolegend) in FACS buffer (5% FBS in PBS) for 15 minutes at 4°C in dark. 100μl FACS buffer was added into the tubes and centrifuged at 500 x g for 5 minutes. Cells were washed with 150μl FACS buffer and centrifuged at 500 x g for 5 minutes twice. The pellet was resuspended in 200μl FACS buffer and run with LSRII flow cytometer (Biolegend). Results were analysed with FlowJo (Treestar).

#### Immunofluorescence

The frozen tumour sections were put at RT and flooded with ice-cold acetone for 10 minutes to fix the tissue. The slides were air-dried and rehydrated in 0.5% Tween-PBS (PBST) for 3 minutes. The non-specific binding of the secondary antibody was blocked with protein block serum-free (Dako X0909) at RT for 30 minutes and incubated with following antibodies at 1:25 dilution: F4/80 Alexa Fluor 488 (Bio-rad); 1:50 dilutions: NOS2 (Abcam), CD3 APC (Tonbo Bioscience), CA9 (Abcam), and TRIB1 (Millipore); 1:100 dilutions: CD31 Alexa Fluor 674 (Biolegend), MR (Abcam), CD4 Alexa Fluor 488 (Biolegend), CD8 PE (Biolegend), IL-15 (Abcam) for 1 hour at RT. The samples were washed with PBST for 5 minutes twice and incubated with secondary antibody Goat anti-Rabbit IgG (H&L) Dylight 550 (ImmunoReagents) at 1:50 dilution for 1 hour at RT. Slides were washed with PBST for 5 minutes three times and mounted with Antifade mounting medium with DAPI (Life Technology). Slides were kept in the dark overnight at RT and imaged immediately or stored at 4°C. Random areas of 4-5 images were captured using a Leica AF6000 microscope, or Nikon A1 confocal microscope and cells were manually quantified with ImageJ (NIH)

## Supporting information

Combined Supplementary info

## Acknowledgement

The work was supported extensively by the University of Sheffield Biological Services Team and Fiona Wright in the Histology Facility. We would like to thank Dr Yang Li and Dr Benjamin Durham for recruitment of blood from healthy participants Dr Victoria Ridger for overseeing human ethics. We thank Dr Penelope Ottowell for providing the breast cancer cells, and Dr Haider Al-Janabi for providing mammary tumour tissues developed in C57BL/6.

## Author Contributions

Conceptualisation of the study, A.O., G.V., M.M., and E.K-T.; Performing the experiments and data analysis: T.K., J.J., F.J.C.F, S.H, S.C.L., A.H.G, G.V., M.M. and E.K-T; Writing and editing the manuscript: all authors

