## Supplementary material for "TRIB1 regulates tumour growth via controlling tumour-associated macrophage phenotypes and is associated with breast cancer survival and treatment response": Combined Supplementary info

**Supplementary items:**

**The Cardiogenics Consortium**

François Cambien[1], Panos Deloukas[2], Jeanette Eardman[3], Alison H Goodall[4,5], Christian Hengstenberg[6], Willem H Ouwehand[2,7], Nilesh J Samani[4,5], Heribert Schunkert[8]

1 INSERM UMRS 937, Pierre and Marie Curie University (UPMC, Paris 6) and Medical School, 91 Bd de l'Hôpital 75013, Paris, France

2The Wellcome Trust Sanger Institute, Wellcome Trust Genome Campus, Hinxton, Cambridge CB10 1SA, UK

3Medizinische Klinik 2, Universität zu Lübeck, Lübeck Germany;

4Department of Cardiovascular Sciences, University of Leicester, Glenfield Hospital, Groby Road, Leicester, LE3 9QP, UK

5Leicester NIHR Biomedical Research Centre , Glenfield Hospital, Leicester, LE3 9QP, UK

6Klinik und Poliklinik für Innere Medizin II, Universität Regensburg, Germany;

7Department of Haematology, University of Cambridge, Long Road, Cambridge, CB2 2PT, UK and National Health Service Blood and Transplant, Cambridge Centre, Long Road, Cambridge, CB2 2PT, UK

8German Center for Cardiovascular Research, Munich Heart Alliance, D-80636 Munich, Germany.

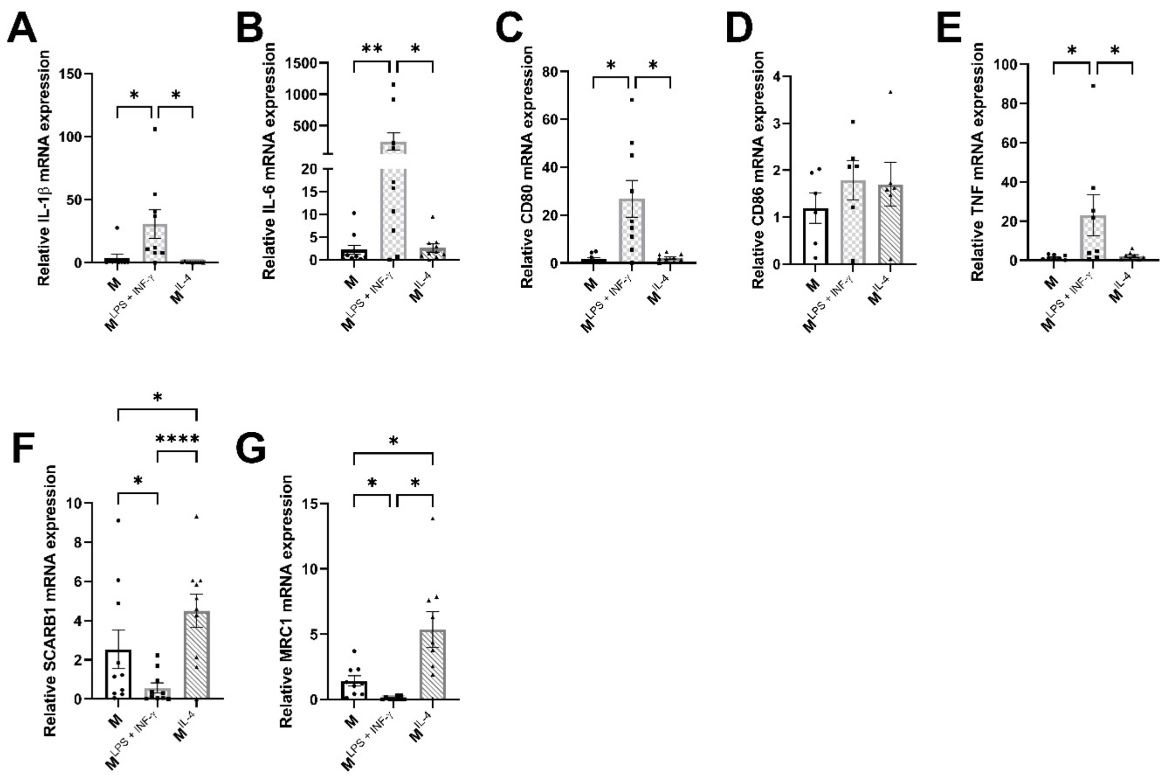


**Supplementary figure 1. Classification of macrophage phenotypes with RT-qPCR.** Human MDMs isolated and differentiated from blood were unstimulated (M) or stimulated with LPS and INF-γ ($M1^{LPS+INF\text{-}\gamma}$) and IL-4 ($M2a^{IL\text{-}4}$) for 24 hours. RNA levels of *IL-1β*, *IL-6, CD80, CD86, TNF, SCARB1, and MRC1* 24 hours after macrophage polarisation. Results of one-way ANOVA are presented; mean±SEM is plotted; *p<0.05 **p<0.01 ****p<0.0001 (n=6-10 donor/group).

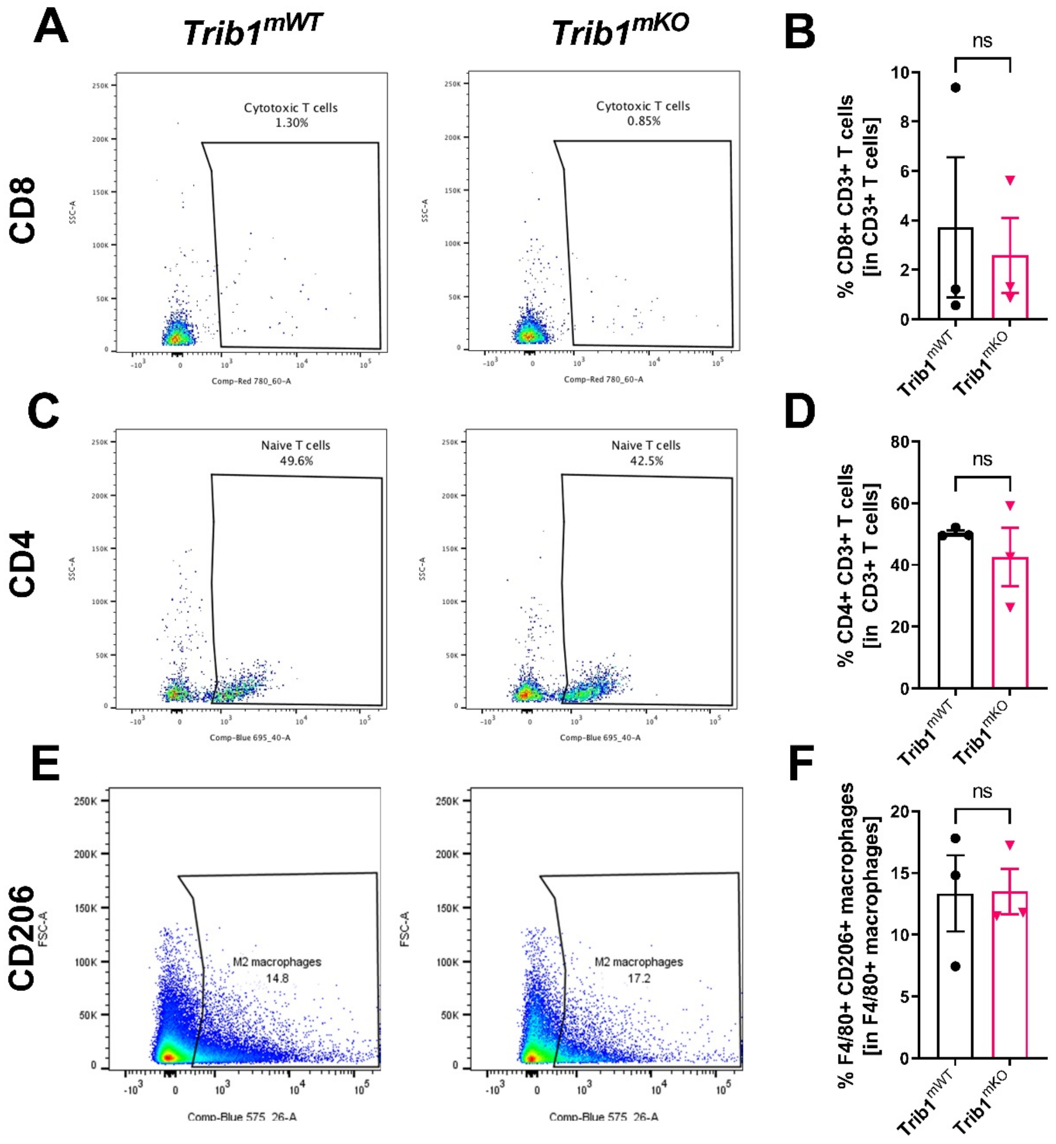


**Supplementary figure 2. *Trib1$^{mKO}$* tumours did not alter T cell subtypes and TAM phenotype. (A, C, E)** Post-mortem analysis of MR+ TAMs in *Trib1$^{mKO}$* and respective *Trib1$^{mWT}$* tumours by flow cytometry. LSRII flow cytometer was used to acquire data and analysed using Flowjo. **(B, D, F)** Quantification of CD4+ naïve and CD8+ cytotoxic T-cellsand MR+ anti-inflammatory TAMs in *Trib1$^{mTg}$* and respective *Trib1$^{mWT}$* tumours. Results of unpaired t-test are presented; mean±SEM is plotted (n=3 mice/group).

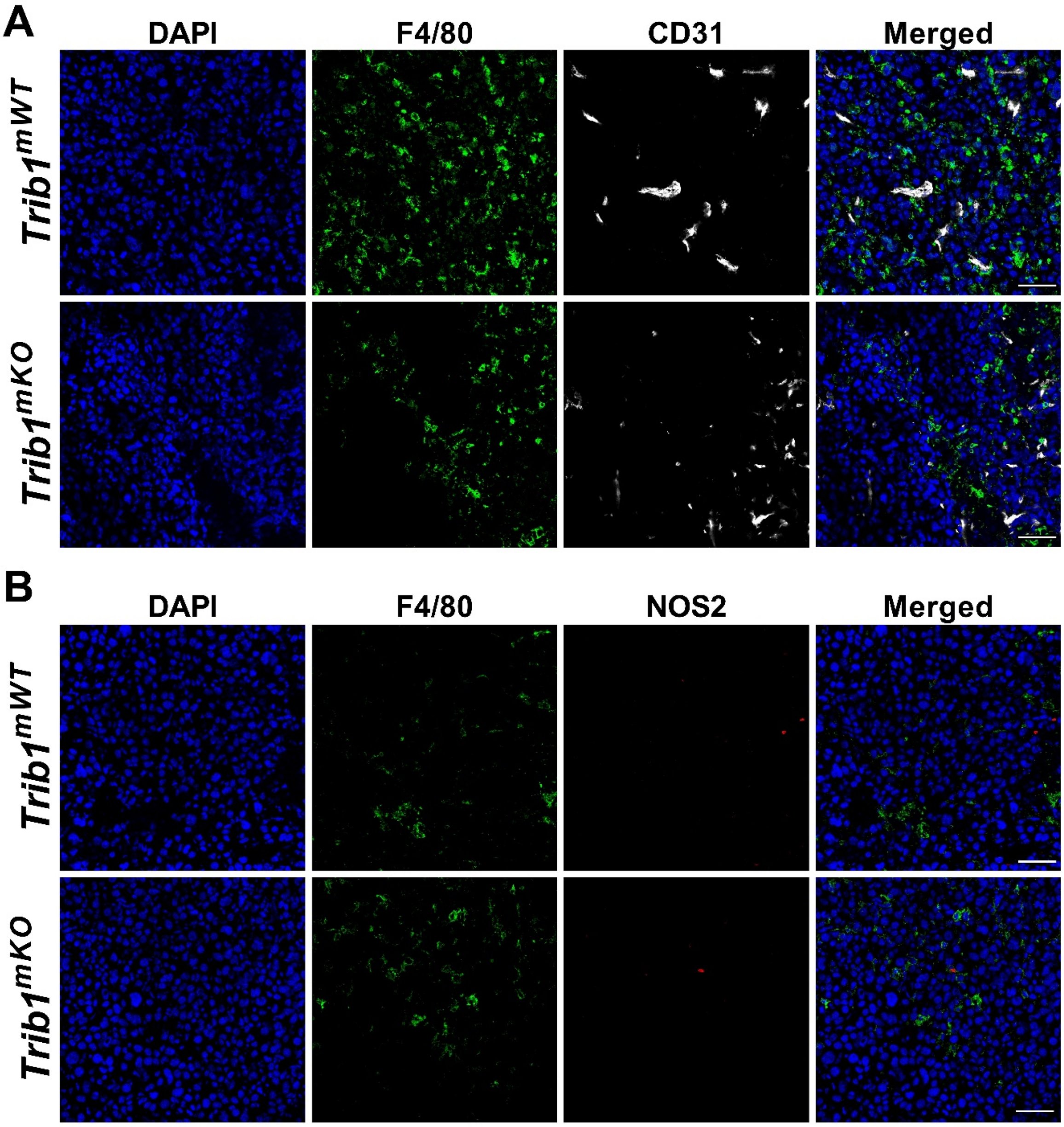


**Supplementary figure 3. Images of fluorescence staining on *Trib1*$^{mKO}$ tumours. (A)** Representative images of CD31 (white) and F4/80 (green) fluorescence staining in *Trib1*$^{mKO}$ and respective *Trib1*$^{mWT}$ tumours (Scale: 50µm). **(B)** Representative images of NOS2 (red) and F4/80 (green) fluorescence staining in *Trib1*$^{mKO}$ and respective *Trib1*$^{mWT}$ tumours (Scale: 50µm). Images were captured using Nikon A1 confocal microscope.

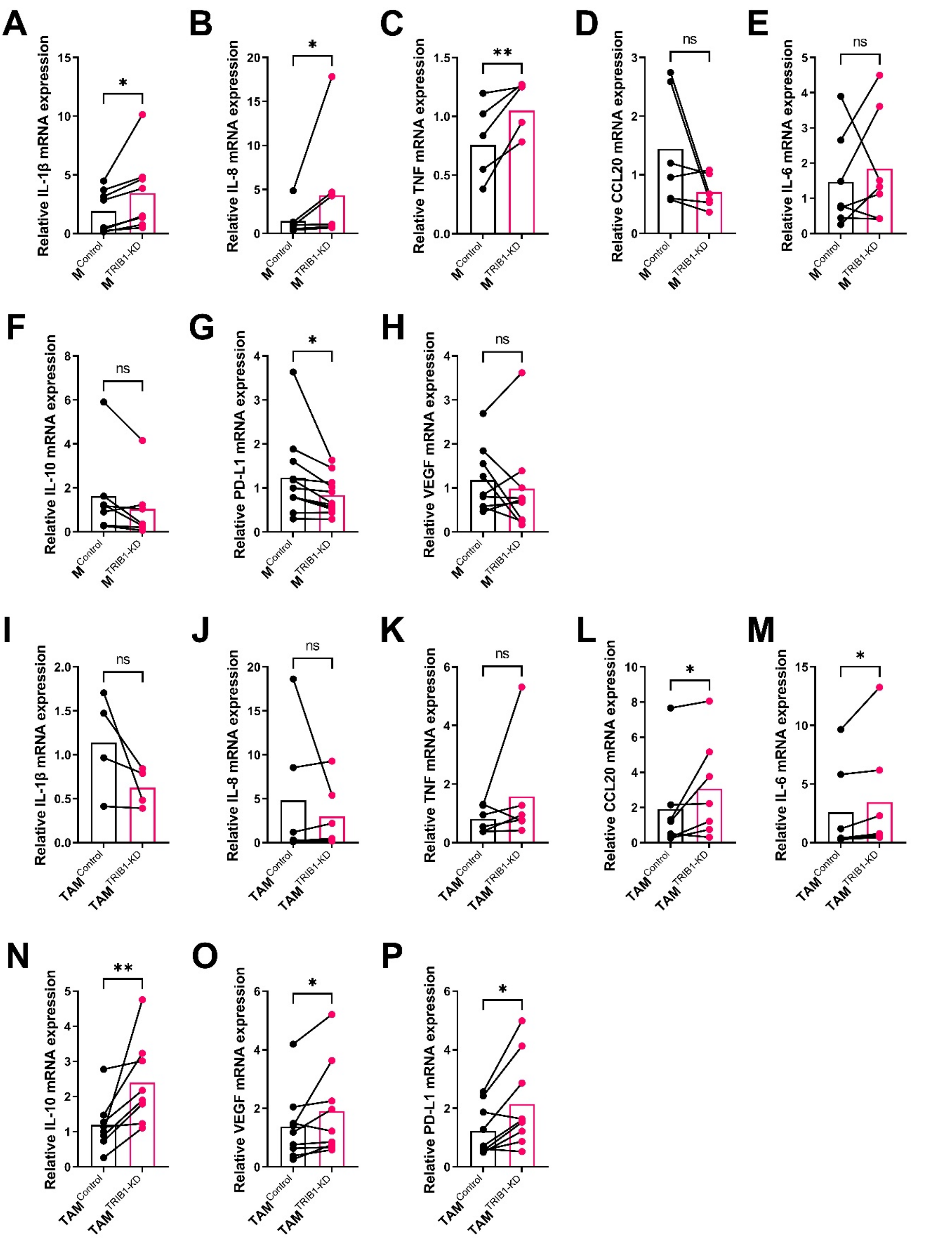


**Supplementary figure 4. *TRIB1* knockdown MDMs accelerate pro-inflammatory cytokines. (A-H)** *In vitro* assessment of cytokine expressions in human MDMs 48 hours after *TRIB1* siRNA transfection. RNA levels of *IL-1β*, *IL-8, TNF, CCL20*, *IL-6, IL-10*, *PD-L1*, and *VEGF* 48 hours after *TRIB1* siRNA transfection ($M^{TRIB1\text{-}KD}$). Results of paired t-test is presented; mean is plotted; *p<0.05 **p<0.01 (n=6-9 donor/group). **(I-P)** *In vitro* assessment of cytokine expressions in human MDMs 72 hours after *TRIB1* siRNA transfection and polarisation towards TAM with CM. RNA levels of *IL-1β*, *IL-8, TNF, CCL20*, *IL-6, IL-10*, *PD-L1*, and *VEGF* 48 hours after *TRIB1* siRNA transfection and TAM polarisation ($TAM^{TRIB1\text{-}KD}$). Results of paired t-test is presented; mean is plotted; *p<0.05 **p<0.01 (n=4-9 donor/group).

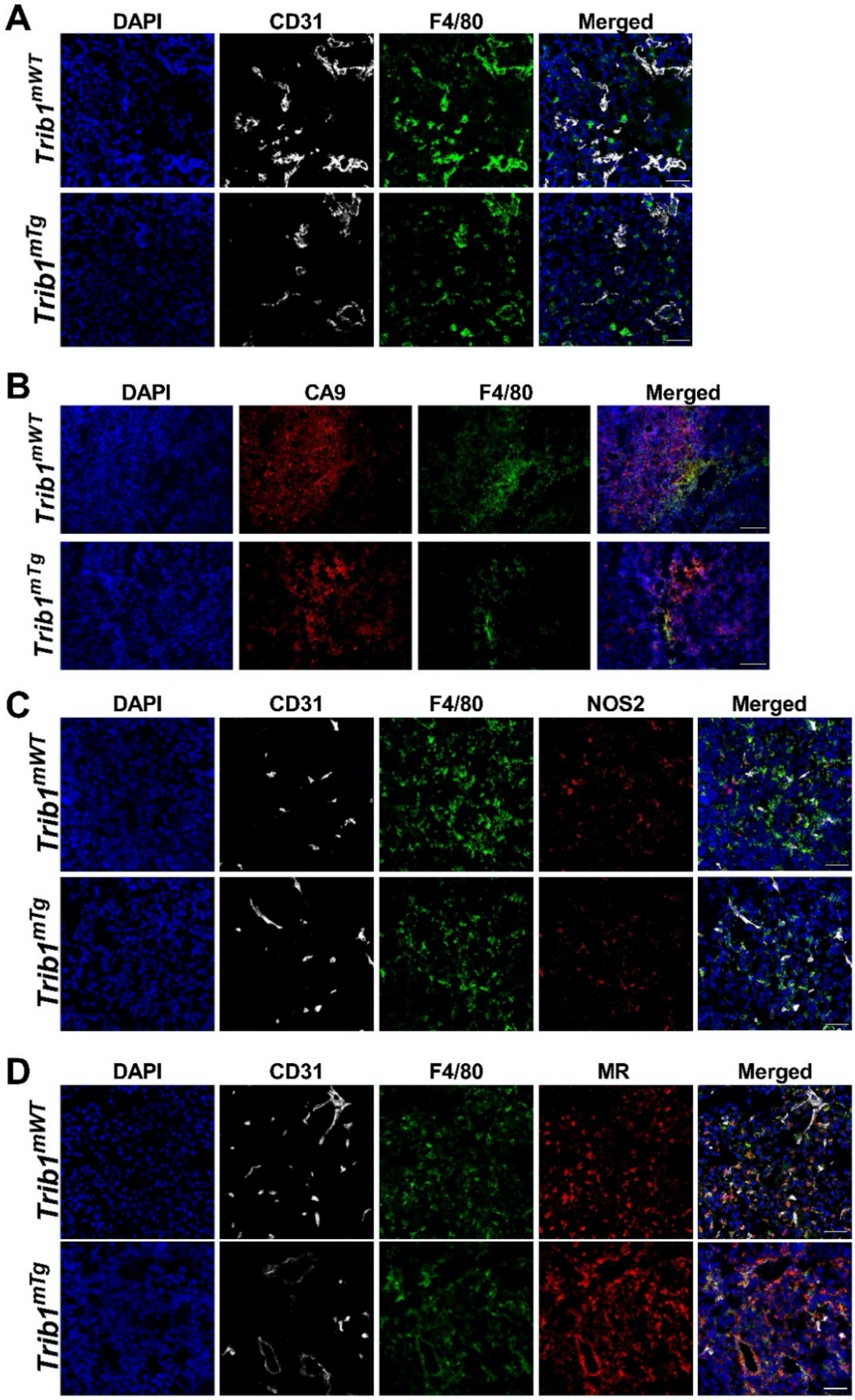


**Supplementary figure 5. Fluorescence staining images of TAMs in *Trib1$^{mTg}$* tumours.** (A) Representative images of CD31 (white) and F4/80 (green) fluorescence staining in *Trib1$^{mTg}$* and respective *Trib1$^{mWT}$* tumours (Scale: 50μm). Images were captured using Nikon A1 confocal microscope. (B) Representative images of CA9 (red) and F4/80 (green) fluorescence staining in *Trib1$^{mTg}$* and respective *Trib1$^{mWT}$* tumours (Scale: 100μm). Images were captured using Leica AF6000 microscope. (C) Representative images of CD31 (white), NOS2 (red) and F4/80 (green) fluorescence staining in *Trib1$^{mTg}$* and respective *Trib1$^{mWT}$* tumours (Scale: 50μm). (D) Representative images of CD31 (white), MR (red) and F4/80 (green) fluorescence staining in *Trib1$^{mTg}$* and respective *Trib1$^{mWT}$* tumours (Scale: 50μm). Images were captured using Nikon A1 confocal microscope.

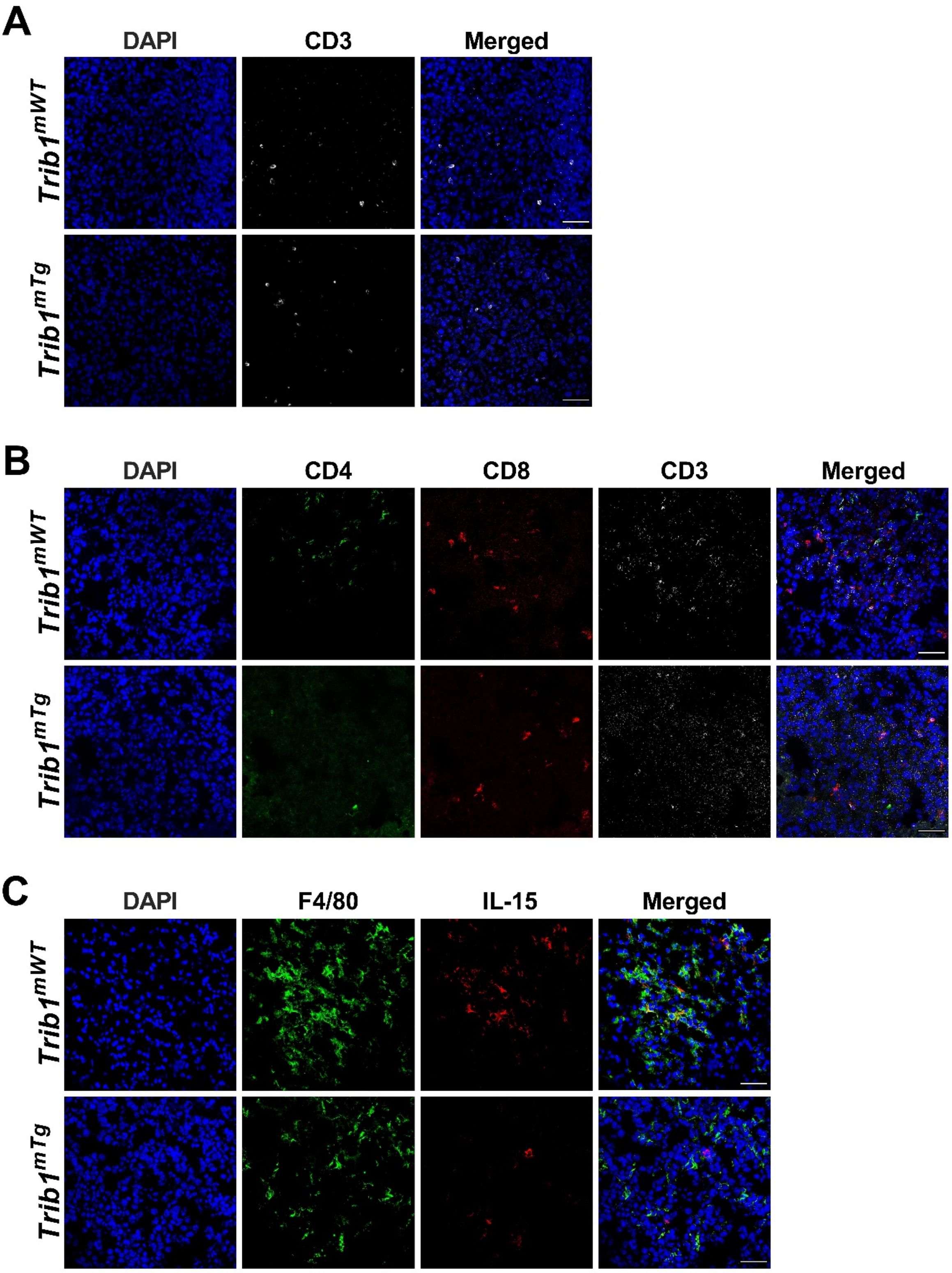


**Supplementary figure 6. Fluorescence staining images of T-cellsand IL-15 expression in TAMs in *Trib1$^{mTg}$* tumours.** (A) Representative images of CD3 (white) fluorescence staining in *Trib1$^{mTg}$* and respective *Trib1$^{mWT}$* tumours (Scale: 50µm). (B) Representative images of CD4 (green), CD8 (red) and CD3 (white) fluorescence staining in *Trib1$^{mTg}$* and respective *Trib1$^{mWT}$* tumours (Scale: 50µm). (C) Representative images of IL-15 (red) and F4/80 (green) fluorescence staining in *Trib1$^{mTg}$* and respective *Trib1$^{mWT}$* tumours (Scale: 50µm). Images were captured using Nikon A1 confocal microscope.

**Supplementary table 1. SYBR RT-qPCR primer sequences.**

| Gene | Species | Forward primer 5’ – 3’ | Reverse primer 5’ – 3’ |
|---|---|---|---|
| **IL-1β** | Human | GCTCGCCAGTGAAATGATGG | GAAGCCCTTGCTGTAGTGGT |
| **IL-6** | Human | ACCCCCAGGAGAAGATTCCA | GATGCCGTCGAGGATGTACC |
| **IL-8** | Human | TGCCAAGGAGTGCTAAAG | CTCCACAACCCTCTGCAC |
| **IL-10** | Human | GCCTTTAATAAGCTCCAAGAG | ATCTTCATTGTCATGTAGGC |
| **IL-15** | Human | ACAGAAGCCAACTGGGTGAA | GCTGTTACTTTGCAACTGGGG |
| **SCARB1** | Human | GAATCCCCATGAACTGCTCTGT | TCCCAGTTTGTCCAATGCCTG |
| **MRC1** | Human | AGATGGGTGGGTTATTTACAAAGA | ATATTTCCATAGAAACTTCTTTTCACTT |
| **TNF** | Human | CCTGCTGCACTTTGGAGTGA | CTTGTCACTCGGGGTTCGAG |
| **PD-L1** | Human | AGGGCATTCCAGAAAGATGAGG | GGTCCTTGGGAACCGTGAC |
| **VEGF** | Human | ATGCGGATCAAACCTCACCA | GCTCTATCTTTCTTTGGTCTGC |
| **CCL20** | Human | ACTGGGTACTCAACACTGAGC | CAAAGCAGCCAGGAGCAAAC |
| **TRIB1** | Human | CTCCACGGAGGAGAGAACCC | GACAAAGCATCATCTTCCCCC |
| **GAPDH** | Human | ATTGCCCTCAACGACCACTTT | CCCTGTTGCTGTAGCCAAATTC |
| **IL-15** | Mouse | GACACCACTTTATACACTGACAGTG | TCACATTCCTTGCAGCCAGA |
| **B-actin** | Mouse | GGGACCTGACAGACTACCTCATG | GTCACGCACGATTTCCCTCTCAGC |

**Supplementary table 2. TRIB1 affected top 10 macrophage pathway analysis.**

| Pathway | Monocyte | | Macrophage | |
|---|---|---|---|---|
| | log.fold.change | FDR | log.fold.change | FDR |
| **CREATION OF C4 AND C2 ACTIVATORS** | | | 0.344 | 0.000073 |
| **TRANSLOCATION OF ZAP 70 TO IMMUNOLOGICAL SYNAPSE** | -0.03364 | 0.536811 | 0.202 | <0.0000001 |
| **PD1 SIGNALLING** | -0.01899 | 0.670612 | 0.172 | <0.0000001 |
| **PHOSPHORYLATION OF CD3 AND TCR ZETA CHAINS** | -0.02801 | 0.544339 | 0.161 | <0.0000001 |
| **HDL MEDIATED LIPID TRANSPORT** | 0.008707 | 0.434654 | 0.159 | <0.0000001 |
| **CHEMOKINE RECEPTORS BIND CHEMOKINES** | 0.228 | <0.0000001 | 0.15 | <0.0000001 |
| **INITIAL TRIGGERING OF COMPLEMENT** | 0.007888 | 0.7942 | 0.149 | 0.0017 |
| **GENERATION OF SECOND MESSENGER MOLECULES** | -0.01264 | 0.668659 | 0.127 | <0.0000001 |
| **LIPOPROTEIN METABOLISM** | | | 0.112 | <0.0000001 |
| **NOREPINEPHRINE NEUROTRANSMITTER RELEASE CYCLE** | 0.04296 | 0.002402 | 0.095 | 0.0000095 |
